# Functional coupling between TRPV4 channel and TMEM16F modulates human trophoblast fusion

**DOI:** 10.1101/2021.12.11.472241

**Authors:** Yang Zhang, Pengfei Liang, Ke Zoe Shan, Liping Feng, Yong Chen, Wolfgang Liedtke, Huanghe Yang

**Affiliations:** Department of Biochemistry, Duke University Medical Center, Durham, NC 27710, USA; Department of Obstetrics and Gynecology, Duke University Medical Center, Durham, NC, 27710, USA; MOE-Shanghai Key Laboratory of Children’s Environmental Health, Xinhua Hospital, Jiao; Department of Neurology, Duke University Medical Center, Durham, NC 27710, USA; Department of Neurobiology, Duke University Medical Center, Durham, NC 27710, USA; Department of Anesthesiology, Duke University Medical Center, Durham, NC 27710, USA; College of Dentistry, Department of Molecular Pathobiology, NYU, New York, NY 10021; Global Development Scientific Council, Regeneron Pharmaceuticals, Tarrytwon NY 10591

## Abstract

TMEM16F, a Ca^2+^-activated phospholipid scramblase (CaPLSase), is critical for placental trophoblast syncytialization, HIV infection, and SARS-CoV2-mediated syncytialization. How TMEM16F is activated during cell fusion is unclear. Here, we used trophoblasts as a model for cell fusion and demonstrated that Ca^2+^ influx through Ca^2+^ permeable transient receptor potential vanilloid channel TRPV4 is critical for TMEM16F activation and subsequent human trophoblast fusion. GSK1016790A, a TRPV4 specific agonist, robustly activates TMEM16F in trophoblasts. Patch-clamp electrophysiology demonstrated that TRPV4 and TMEM16F are functionally coupled within Ca^2+^ microdomains in human trophoblasts. Pharmacological inhibition or gene silencing of TRPV4 hindered TMEM16F activation and subsequent trophoblast syncytialization. Our study uncovers the functional expression of TRPV4 and a physiological activation mechanism of TMEM16F in human trophoblasts, thus providing us with novel strategies to regulate CaPLSase activity as a critical checkpoint of physiologically- and disease-relevant cell fusion events.

## Introduction

Phospholipids as essential building blocks of mammalian cell membranes are asymmetrically distributed (1, 2). TMEM16F Ca^2+^-activated phospholipid scramblase (CaPLSase), is a passive phospholipid transporter that catalyzes trans-bilayer movement of the phospholipids down their concentration gradients (3, 4). In response to intracellular Ca^2+^ increase, phosphatidylserine (PS), an anionic phospholipid always residing in the inner membrane leaflet, can rapidly permeate through a hydrophilic pathway of TMEM16F and become exposed to the cell surface (5–8). In platelets, TMEM16F CaPLSase-mediated PS exposure creates a pro-coagulant surface to facilitate thrombin generation and blood coagulation (1, 3, 9). Consistent with its important role in blood coagulation, TMEM16F deficiency leads to deficient thrombin generation and prolonged bleeding in Scott syndrome patients (3, 10, 11) and in TMEM16F knockout mice (9). Besides blood coagulation, TMEM16F-mediated PS exposure has been increasingly reported to be important in cell-cell fusion and viral-cell fusion events including placental trophoblast syncytialization (12), SARS-CoV2-mediated syncytialization (13) and HIV infection (14). In order to elucidate the physiological and pathological roles of TMEM16F in these cell-cell and viral-cell fusion events, one hitherto missing prerequisite is to understand the cellular mechanisms of TMEM16F activation.

Ca^2+^ binding to the highly conserved Ca^2+^ binding sites is required for the activation of the TMEM16 family of proteins (15–18). However, the apparent Ca^2+^ sensitivity of TMEM16F is lower than that of TMEM16A and TMEM16B, which are Ca^2+^-activated Cl^-^ channels (CaCCs) (4, 9, 18). Different from the CaCCs that can be readily activated by both Ca^2+^ entry through Ca^2+^ channels and Ca^2+^ release from internal stores (9, 19–24), TMEM16F activation usually requires more robust and sustained intracellular Ca^2+^ elevation (3, 4, 8, 9, 25-29). To induce higher intracellular Ca^2+^ levels, Ca^2+^ ionophores have been widely used to artificially activate TMEM16F and study its biophysical properties or its functional expression in native cells (3, 5, 8, 12). Nevertheless, how TMEM16F is activated under physiological conditions has remained unclear, which greatly hinders improved understanding of TMEM16F biology and subsequent rationally-based modulation of its activity.

We recently showed that TMEM16F is highly expressed in placenta trophoblasts and plays an essential role in human trophoblast syncytialization *in vitro* and mouse placental development *in vivo* (12). Our TMEM16F gene-targeted mice displayed deficiency in trophoblast syncytialization, signs of intrauterine growth restriction, and partial perinatal lethality. Our animal experiments suggested that the extremely rare incidence of Scott syndrome with only mild bleeding tendency (30, 31) could be derived from pregnancy complications due to *TMEM16F* loss-of-function in trophoblasts. In order to further understand the underlying mechanism of TMEM16F-mediated trophoblast fusion, it is critical to elucidate how TMEM16F is activated under physiological conditions. Ca^2+^-permeable TRPV6 and L-type voltage-gated Ca^2+^ (Ca_V_) channels have been reported to be expressed in trophoblasts (32–36). Due to its fast inactivation kinetics upon Ca^2+^ entry (37), TRPV6 presumably is involved in trans-placental Ca^2+^ transport (CaT) instead of sustained Ca^2+^ signaling (38, 39). L-type Cav channel, on the other hand, requires strong membrane depolarization (40), which is hardly achieved in non-excitable trophoblasts. Therefore, these known trophoblast Ca^2+^ channels are unlikely to play a major role in activating TMEM16F during trophoblast fusion.

Here we report that Ca^2+^ influx through Ca^2+^-permeable TRPV4 channels in human trophoblasts plays an important role in activating TMEM16F CaPLSase and modulating trophoblast fusion. Using Ca^2+^ imaging and patch clamping, we showed that TRPV4 channel is functionally expressed in human trophoblasts. GSK-1016790A (GSK101), a TRPV4 specific agonist, triggered robust Ca^2+^ entry, which stimulated TMEM16F CaPLSase activation and subsequent PS surface exposure. Utilizing TMEM16F’s ion channel function and different Ca^2+^ chelating kinetics of EGTA and BAPTA, our patch clamp recording further demonstrated that trophoblast TPRV4 and TMEM16F are functionally coupled within microdomains. We also demonstrated that gene silencing of TRPV4 or pharmacological inhibition of TRPV4 channels hindered trophoblast syncytialization. Therefore, manipulating Ca^2+^ channels can serve as an effective approach to control TMEM16F activities, thereby helping dissect the biological functions of the CaPLSase. We anticipate that our findings will advance the understanding of the cellular activation mechanisms of TMEM16 CaPLSases in different physiological and pathological fusion processes, and also shine light on devising new rationally-based therapeutic strategies to target TMEM16F-mediated diseases including HIV infection and COVID19.

## Results

### TRPV4 is functionally expressed in human trophoblasts

As a contiguous multinucleated cell layer in the chorionic villi, placental syncytiotrophoblasts establish the most important materno-fetal exchange interface and perform vital barrier, transport, and secretion functions to sustain fetal development and maternal health (Fig. 1A) (41, 42). A syncytiotrophoblast is formed and maintained by continuous cell fusion of underlying mononucleated cytotrophoblasts into the syncytial layer during placental development (42). Besides the trophoblast CaT encoded by *TRPV6* (38, 39), the single cell RNA sequencing results from the Human Protein Atlas showed that *TRPV4* is also highly expressed in cytotrophoblasts and syncytiotrophoblasts in the human placenta among the TRPV channels (Fig. S1) (43, 44). Consistent with the database, our RT-qPCR experiment also showed that TRPV4 transcripts are expressed in human primary trophoblasts and the human choriocarcinoma trophoblast cell line BeWo (Fig. 1B, 1C). Our immunofluorescence using a specific anti-TRPV4 antibody (Fig. S2) further demonstrated that TRPV4 proteins are expressed in the multi-nucleated syncytiotrophoblasts and single-nucleated cytotrophoblasts in human placenta villi, as well as on the BeWo cell surface (Fig. 1D, 1E).

**Figure 1.**
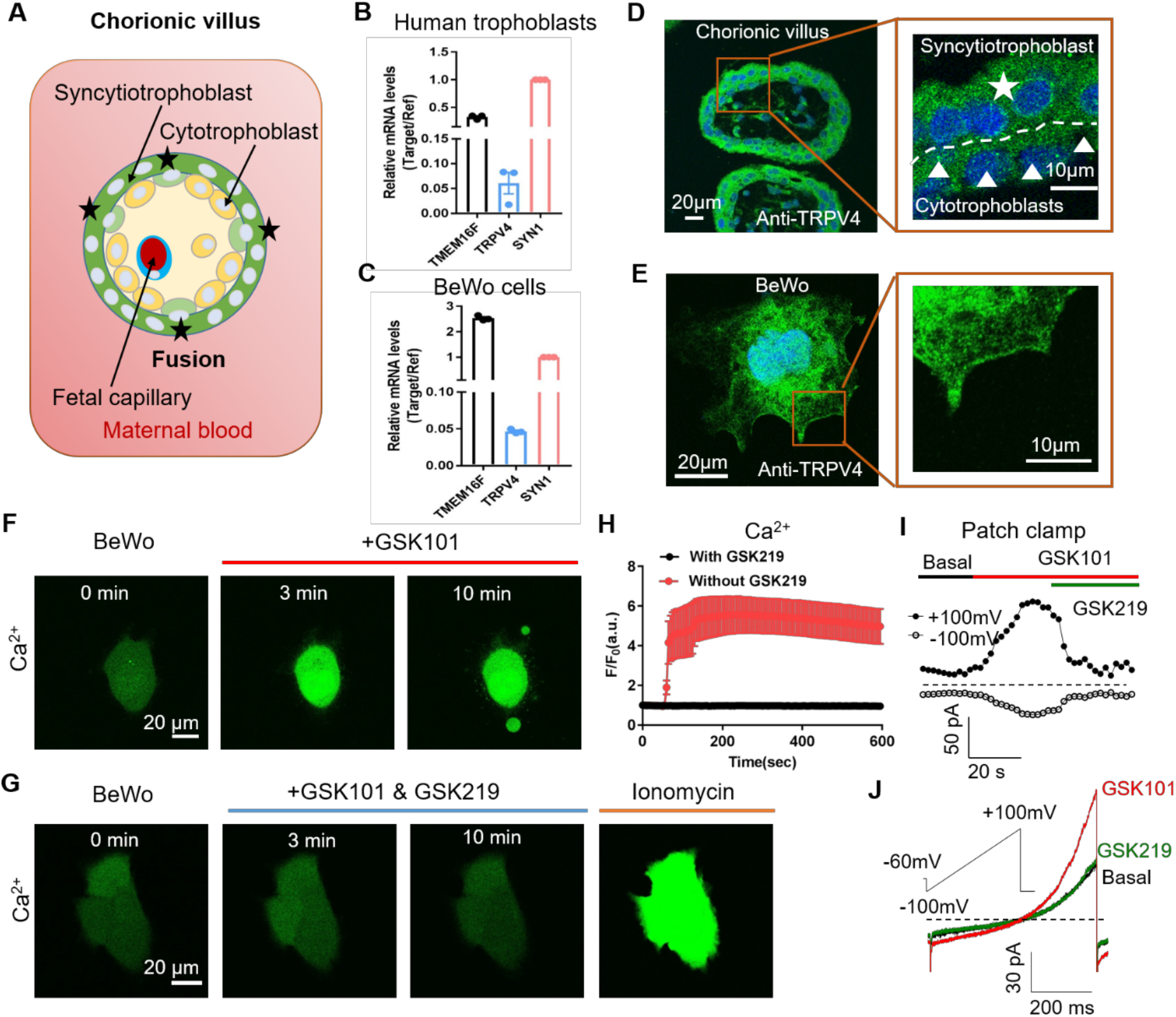
TRPV4 is functionally expressed in human trophoblasts. (**A**) Schematic of the first trimester placental villus and trophoblast fusion. **(B-C)** qRT-PCR of *TRPV4* in primary human trophoblasts (B) and BeWo cells (C). All genes were normalized to GAPDH and then normalized to Syncytin-1 (SYN1). **(D)** Representative immunofluorescence of TRPV4 (green) and nuclei (blue) in a human first trimester placenta villus (cross-section). TRPV4 is expressed in both cytotrophoblasts and syncytiotrophoblasts. **(E)** Representative immunofluorescence of TRPV4 (green) in BeWo cells. The nuclei were stained with Hoechst (blue). All fluorescence images are the representatives of at least three biological replicates. **(F)** GSK1016790A (GSK101, 20 nM), a specific TRPV4 agonist, triggered robust intracellular Ca^2+^ elevation in BeWo cells. **(G)** GSK2193874 (GSK219, 500 nM), a selective TRPV4 antagonist, completely prevented GSK101-induced Ca^2+^ influx through TRPV4 channels in BeWo cells. Ca^2+^ elevation through ionomycin was intact in the presence of GSK219. **(H)** Summary of GSK101 and GSK219 effects on BeWo cell Ca^2+^ dynamics measured by Ca^2+^ dye (Calbryte 520). All fluorescence images are the representatives of at least three biological replicates. **(I-J)** Time course of outside-out currents elicited in response to 30 nM GSK101 with and without 500 nM GSK219 (I) and representative current traces in Panel I in the presence of GSK101 and GSK219+GSK101 (J). The currents were elicited by a 500 ms ramp voltage protocol from −100 mV to +100 mV. The holding potential was set at −60 mV.

TRPV4 channels is a Ca^2+^ permeable, non-selective cation channel with much less pronounced inactivation than TRPV6 (37, 45, 46). Its activation thus can lead to sustained Ca^2+^ elevation in trophoblasts, which would subsequently activate TMEM16F. The availability of selective and potent TRPV4 agonists such as GSK101 and antagonists including GSK-2193874 (GSK219) (47, 48) enabled us to explicitly confirm TRPV4’s functional expression in human trophoblasts and examine its capability to activate TMEM16F. We found that 20 nM GSK101 can robustly increase intracellular Ca^2+^ in BeWo cells, which sustained for no less than 10 minutes (Fig. 1F, 1H). Application of 20 µM 4α-phorbol-12,13-didecanoate (4α-PDD), another TRPV4 agonist, also induced intracellular Ca^2+^ increase in BeWo cells (Fig. S3A). On the other hand, 500 nM TRPV4 antagonist GSK219 completely prevented 20 nM GSK101-induced intracellular Ca^2+^ increase, yet it failed to inhibit ionomycin-induced Ca^2+^ increase (Fig. 1G–H). Our Ca^2+^ imaging experiments using specific pharmacological tools thus support the concept that TRPV4 is functionally expressed in human trophoblasts.

To further validate our Ca^2+^ imaging results, we used outside-out patch clamp to record GSK101-induced current from BeWo cell membranes. Indeed, GSK101 robustly elicited an outward rectifying cation current that could be rapidly inhibited by co-application of GSK219 (Fig. 1I–J). Consistent with the Ca^2+^ imaging experiments (Fig. 1F), our electrophysiological experiments further support that TRPV4 channel is functionally expressed in human trophoblasts. Consistent with results using TRPV4 antagonist GSK219 (Fig. 1G–H), silencing TRPV4 with siRNAs also greatly reduced GSK101-induced Ca^2+^ elevation (Fig. S3B) and the non-selective, outward rectifying current (Fig. S3C-E). Taken together, our Ca^2+^ imaging and patch clamp results established that Ca^2+^ permeable TRPV4 channel is functionally expressed in human trophoblasts and that TRPV4 activation can lead to sustained Ca^2+^ elevation in these cells.

### Ca^2+^ entry through TRPV4 activates TMEM16F in human trophoblasts

Utilizing TMEM16F’s dual function as a Ca^2+^-activated (i) phospholipid scramblase and (ii) a non-selective ion channel, we applied a fluorescence imaging-based CaPLSase assay (5, 8, 12) and patch clamp technique to investigate whether TRPV4-mediated Ca^2+^ influx can activate endogenous TMEM16F in human trophoblasts. Upon GSK101 application, Ca^2+^ increased in both BeWo cells (Fig. 2A, 2C) and primary human term placental trophoblasts (Fig. S4), followed by phospholipid scrambling as evidenced by robust cell surface accumulation of the fluorescently tagged PS sensor Annexin-V (AnV) (Fig. 2A, 2D, Video S1). In stark contrast, TMEM16F deficient BeWo (KO) cells did not exhibit any CaPLSase activity, despite robust Ca^2+^ elevation in response to GSK101 stimulation (Fig. 2B, 2C–D, Video S2). Our fluorescence imaging experiments thus indicated that TRPV4 activation can robustly activate TMEM16F CaPLSases in human trophoblasts.

**Figure 2.**
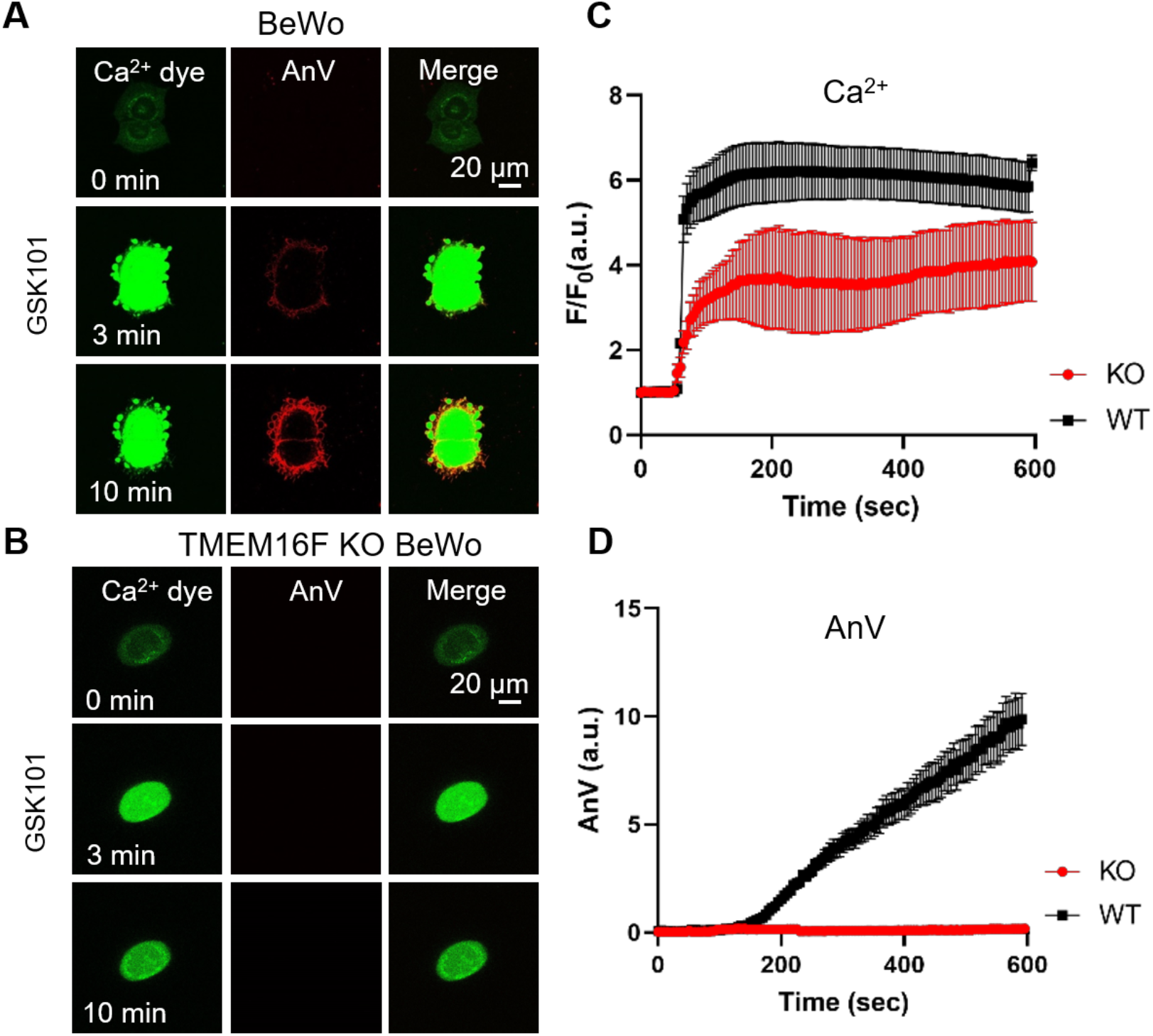
Ca^2+^ influx through TRPV4 activates TMEM16F scramblase. **(A)** 20 nM GSK101 triggered Ca^2+^ influx and PS exposure in BeWo cells. **(B)** 20 nM GSK101 induced Ca^2+^ influx but failed to trigger PS exposure in the TMEM16F knockout (KO) BeWo cell line. Ca^2+^ dye (Calbryte 520, green) and fluorescently tagged AnV proteins (AnV-CF594, red) were used to monitor the dynamics of intracellular Ca^2+^ and PS externalization, respectively. All fluorescence images are the representatives of at least three biological replicates. (**C-D**) Time course of GSK101 triggered Ca^2+^ influx (C) and PS exposure (D) in BeWo WT and TMEM16F KO cells.

We also used patch clamp to record whole cell current of BeWo cells in response to GSK101. GSK101 rapidly triggered an outward rectifying TRPV4 current that plateaued and sustained in both wildtype (WT) and TMEM16F KO BeWo cells (Fig. 3A–C). A much larger, voltage-, and time-dependent conductance developed after a delay of 6-8 minutes (Fig. 3A, D & E). Different from the initial TRPV4 current, the newly evolved conductance showed slow activation and deactivation kinetics (Fig. 3B–C). The biophysical features of the second conductance closely resembled the TMEM16F channels recorded in heterologous expression systems (25–27) and the endogenous TMEM16F channel recorded in WT BeWo cells when infusing 100 µM Ca^2+^ directly into the cytosol via glass pipette (Fig. S5A). In the TMEM16F KO BeWo cells, the Ca^2+^-, voltage- and time-dependent conductance no longer exist (Fig. S5A); and only an outward-rectifying TRPV4-like conductance can be recorded (Fig. 3A–C). Our patch clamp experiments therefore demonstrated that TRPV4 activation can robustly activate TMEM16F channels in human trophoblasts.

**Figure 3.**
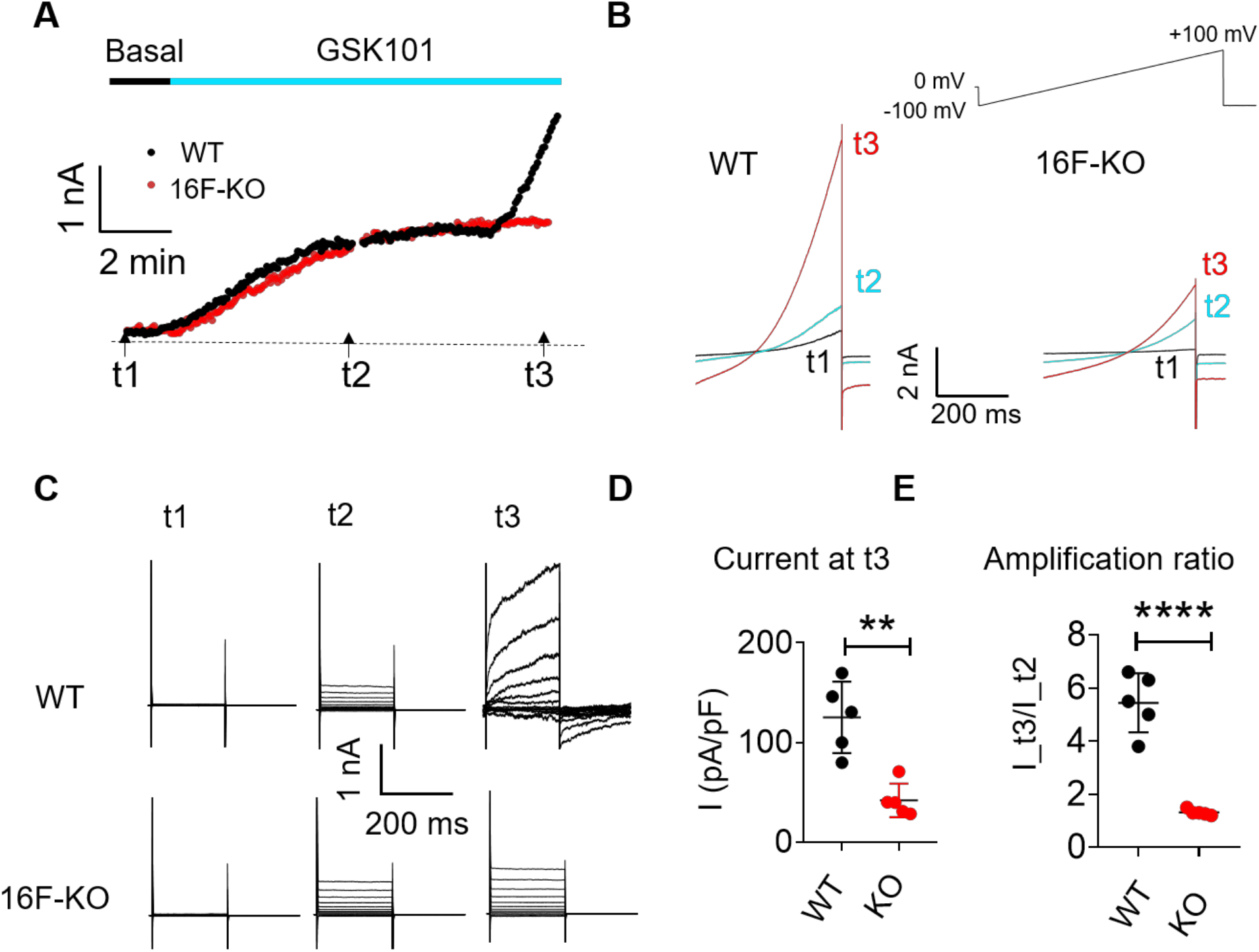
The activation of TRPV4 elicits TMEM16F current. **(A-B)** Time course of whole-cell currents elicited in response to 30 nM GSK101 in WT and TMEM16F-KO BeWo cells. The currents were elicited with a ramp protocol shown in Panel B (top). (A) The current amplitudes at +100 mV were plotted every 5 seconds. (B) Representative currents at three different time points as shown in Panel A. **(C)** Representative current traces elicited by a voltage step protocol (200 ms) from −100 mV to +140 mV at three different time points t1, t2 and t3 as indicated in Panel A. **(D)** Statistical analysis of current density at t3 in WT and TMEM16F-KO BeWo cells. Values represent mean ± SEM and statistics were done using student t-test (n=5 for each group, **: p<0.01). **(E)** Statistical analysis of amplification ratio (current amplitude ratio at t3 and t2) in WT and TMEM16F-KO BeWo cells. Values represent mean ± SEM and statistics were done using student t-test (n=5 for each group, ****: p<0.0001).

Using TRPV4 siRNA and GSK219, we further demonstrated that TRPV4 inhibition robustly suppressed GSK101-induced Ca^2+^ elevation and eliminated subsequent TMEM16F-CaPLSase activation (Fig. 4A–B). Meanwhile, siRNA knockdown of TRPV4 not only greatly suppressed the endogenous TRPV4 current in BeWo cells, but also prevented the development of the voltage-and time-dependent current contributed by TMEM16F (Fig. 4C–E). Our CaPLSase imaging and patch clamp experiments thus demonstrated that TRPV4, a hitherto under-appreciated Ca^2+^-permeable channel in human trophoblasts, can serve as an upstream Ca^2+^-source for trophoblast TMEM16F activation.

**Figure 4.**
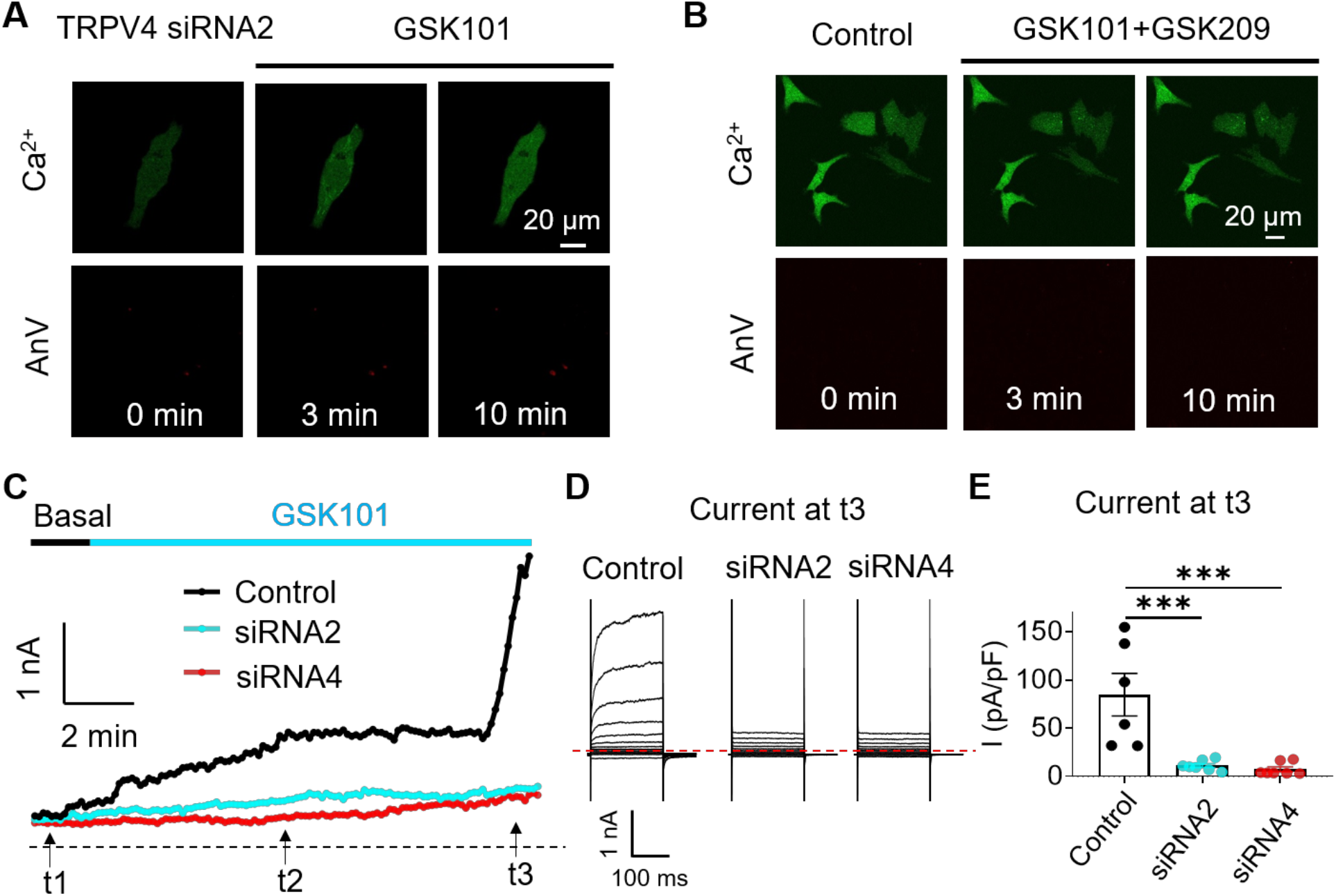
siRNA knockdown of TRPV4 abolishes GSK101-induced Ca^2+^ influx and subsequent TMEM16F activation. **(A)** CaPLSase activity was diminished in TRPV4 knockdown BeWo cells. In BeWo cells with TRPV4 knockdown, fluorescently-tagged AnV, an indicator for externalized PS, and Ca^2+^ dye show no intensity change after the stimulation of GSK101. **(B)** CaPLSase activity and intracellular Ca^2+^ increasing induced by 20 nM GSK101 was eliminated by 500 nM GSK219. **(C)** Time course of whole-cell currents elicited in response to GSK101 in scramble siRNA (control) or TRPV4 siRNAs treated BeWo cells. The currents were elicited with a ramp protocol from −100 mV to +100 mV and plotted every 5 seconds at +100 mV. **(D)** Representative current traces elicited by a voltage step protocol (200 ms) from −100 mV to +140 mV at three different time points t1, t2 and t3 as indicated on panel C. **(E)** Statistical analysis of current density at t3 in WT and TRPV4-siRNA knockdown BeWo cells. Current densities after scrambled siRNA, TRPV4-siRNA2 and TRPV4-siRNA4 treated are 84.72±21.97, 11.12±2.03 and 7.32±2.53 pA/pF, respectively. Values represent mean ± SEM and statistics were done using student t-test (n=5 for each group, ***: p<0.001).

### TRPV4 and TMEM16F are functionally coupled within microdomains

In order to efficiently activate TMEM16F that has relatively low Ca^2+^ sensitivity (8, 9, 25), its Ca^2+^ sources need to be in the vicinity. To test this, we first conducted immunostaining of TRPV4 and TMEM16F in BeWo cells to estimate their membrane co-localization. To circumvent the incompatibility of the available TRPV4 and TMEM16F antibodies, we infected BeWo cells with a lentivirus carrying mCherry-tagged TMEM16F (12) and used a mouse anti-mCherry to detect the heterologously expressed TMEM16F and a rabbit anti-TRPV4 to detect endogenous TRPV4 channels. As shown in Fig. S6, TRPV4 and TMEM16F-mCherry signals were co-localized on BeWo cell surface, suggesting that these two proteins are in close proximity.

To further validate with a functional method, we used patch clamp, a much more quantitative approach, to measure the proximity between the endogenous TMEM16F and TRPV4 in trophoblasts by taking advantage of TMEM16F’s channel activity and the differential Ca^2+^ buffering kinetics of EGTA and BAPTA to distinguish Ca^2+^ micro-from nano-domains of Ca^2+^-permeable channels (49–52). It has been well documented that there is a long delay of TMEM16F channel action under whole cell configuration (Figs. 3A and Fig. S5A) (8, 25, 27, 53). Interestingly, the delayed activation was not observed under excised inside-out patch configuration (9, 53). Even though the detailed mechanism is still unclear, a recent report suggested that actin cytoskeleton might partially contribute to the observed delay (53). To avoid the complication induced by the delay, we conducted outside-out patch clamp recording (Fig. 5A). Under this patch configuration, GSK101 can be directly applied to the extracellular side to activate TRPV4 channels and TMEM16F current can be immediately elicited by TRPV4-mediated Ca^2+^ influx without delay (Fig. S5B). In the presence of GSK101, we varied the duration of a −100 mV pre-pulse to facilitate Ca^2+^ influx (Fig. 5A). In the absence of extracellular Ca^2+^, a small, instantaneous outward rectifying current was recorded from the WT BeWo cells in response to 30 nM GSK101 (Fig. 5B–C, #1). As this current depended on application of GSK101 (Fig. S7A) but not on the duration of the prepulse, it was mediated by TRPV4. Under 2.5 mM extracellular Ca^2+^ and in the presence of 0.2 mM intracellular EGTA from the electrode, GSK101 rapidly elicited a large, time dependent current in WT (Fig. 5B–C, #2) but not in TMEM16F-KO BeWo cells (Fig. S7B), supporting that TMEM16F current is responsible for the GSK101-induced time dependent current. Interestingly, the TMEM16F current amplitudes remained stable regardless of prepulse duration, suggesting that Ca^2+^ influx through TRPV4 channels can readily activate TMEM16F in the vicinity and then rapidly diffuse into the bulk pipette solution without accumulation (Fig. 5B–C, #2). When the intracellular EGTA concentration was increased to 2 mM, the development of TMEM16F current after GSK101 stimulation was drastically delayed, suggesting that higher concentration of EGTA in the electrode can efficiently buffer Ca^2+^ influx through TRPV4 channels and hinder TMEM16F activation (Fig. 5B–C, #3). When the prepulse was prolonged, more Ca^2+^ entered through TRPV4 channels, which went beyond EGTA’s buffering capability within the Ca^2+^ microdomains and allowed Ca^2+^ to activate nearby TMEM16F. In stark contrast, 2 mM BAPTA, which has much faster Ca^2+^ chelating kinetics (50–52), completely ablated the development of TMEM16F current; and only TRPV4 current could be observed under this condition (Fig. 5B–C, #4). Supporting our immuno-colocalization result (Fig. S6), the rapid TMEM16F channel activation in response to TRPV4 channel opening indicate that the two proteins are spatially close to each other in human trophoblasts. The differential effectiveness of BAPTA but not EGTA to abolish GSK101-induced TMEM16F channel activation suggests that TMEM16F and TRPV4 co-signal in a shared microdomain.

**Figure 5.**
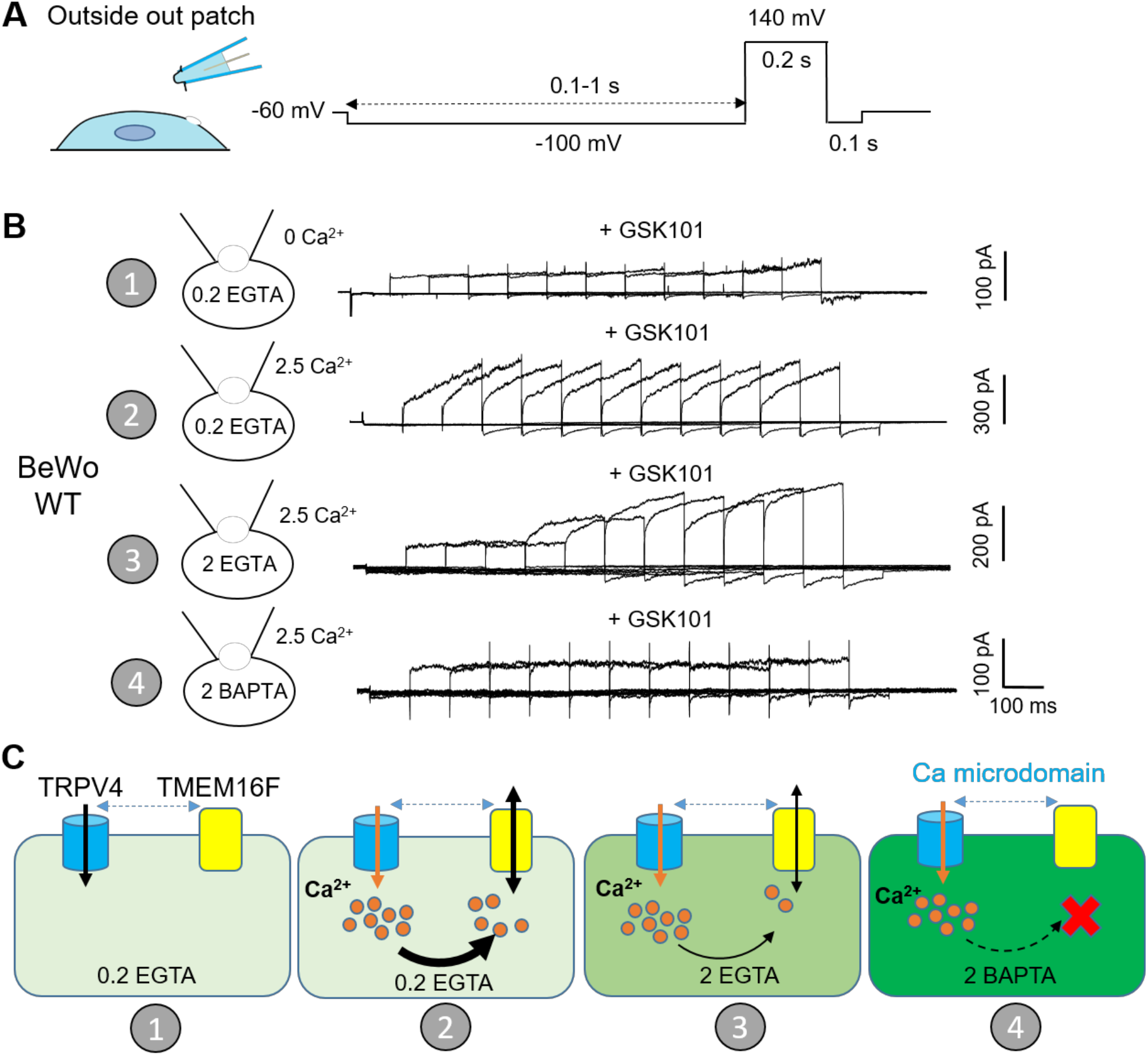
TRPV4 and TMEM16F are functionally coupled within microdomain. **(A)** Outside-out patch configuration and voltage protocols used to demonstrate TRPV4-TMEM16F coupling. The holding potential was set at −60 mV. A −100 mV pre-pulse with varied length from 0.1 second to 1 second was applied along with perfusion of 30 nM GSK101 to induce Ca^2+^ influx. Following the pre-pulse, a depolarized pulse with 0.2 second duration and 140 mV amplitude was applied to record TMEM16F current. **(B)** Representative outside-out patch recordings from wildtype (WT) BeWo cells under four different conditions: 1) intracellular 0.2 mM EGTA, extracellular 0 Ca^2+^ + 30 nM GSK101 (n=5); 2) intracellular 0.2 mM EGTA, extracellular 2.5 mM Ca^2+^ + 30 nM GSK101 (n=5); 3) intracellular 2 mM EGTA, extracellular 2.5 mM Ca^2+^ + 30 nM GSK101 (n=6); 4) intracellular 2 mM BAPTA, extracellular 2.5 mM Ca^2+^ + 30 nM GSK101 (n=6). **(C)** Diagrams to demonstrate TRPV4-TMEM16F coupling under each condition in Panel B. The intensity of green color depicts Ca^2+^ chelating capacity and kinetics.

### Inhibiting TRPV4 hinders trophoblast fusion *in vitro*

We recently showed that TMEM16F plays an indispensable role in controlling trophoblast fusion (12). To elucidate the physiological function of TRPV4-TMEM16F coupling in trophoblasts, we examined the impacts of TRPV4 inhibition on forskolin-induced BeWo cell fusion *in vitro*. Consistent with the critical role of TRPV4 in mediating Ca^2+^ influx and TMEM16F-mediated PS externalization, the fusion index of BeWo cells was significantly decreased when exposed to TRPV4-inhibitor GSK219 (Fig. 6A–B). Extending the results from pharmacological inhibition of TRPV4, TRPV4 siRNA knockdown also significantly hindered BeWo cell fusion (Fig. 6C–D). Our cell-based fusion measurements thus suggested that TRPV4-TMEM16F coupling plays a critical role in controlling trophoblast fusion.

**Figure 6.**
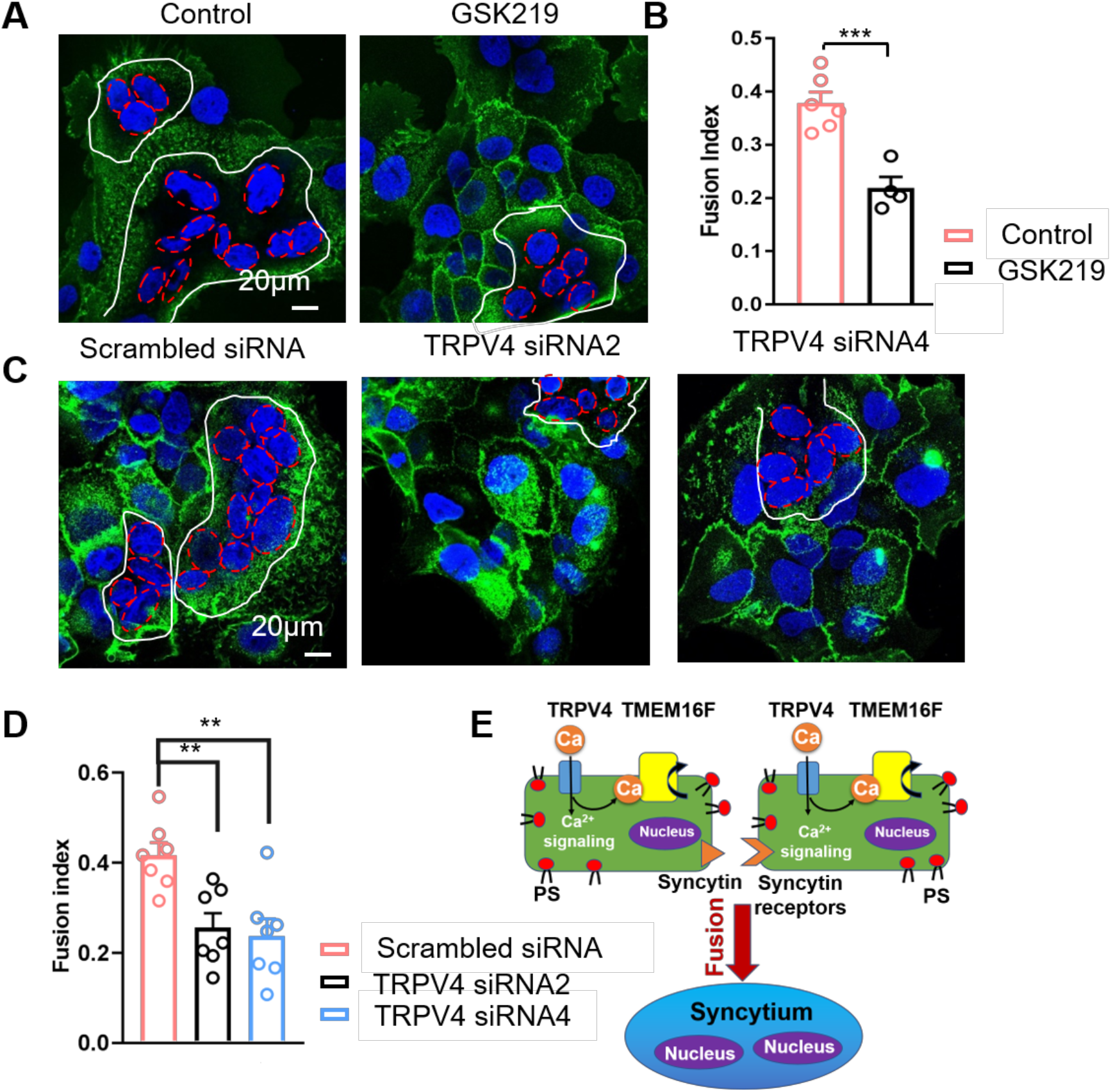
TRPV4 inhibition hinders trophoblast fusion *in vitro* but not *in vivo*. **(A)** Representative images of control and GSK219 treated BeWo cells after 48-hour forskolin treatment. Nuclei and membranes were labelled with Hoechst (blue) and Di-8-ANEPPS (green), respectively. **(B)** GSK219 inhibits forskolin-induced BeWo cell fusion. Unpaired two-sided Student’s *t*-test. ***: p < 0.001, ****: p < 0.0001. Error bars indicate ± SEM. Each dot represents the average of fusion indexes of six random fields from one cover glass. **(C)** Representative images of control (scrambled siRNA) and TRPV4 knockdown (TRPV4 siRNA 2 and 4) BeWo cells after 48-hour forskolin treatment. **(D)** TRPV4 siRNA knockdown inhibits forskolin-induced BeWo cell fusion. Unpaired two-sided Student’s *t*-test. **: p, < 0.01. Error bars represent ± SEM. Each dot represents the average of fusion indexes of six random fields from one cover glass. All fluorescence images are the representatives of at least three biological replicates. **(E)** TRPV4 channel participates in trophoblast Ca^2+^ signaling including activating TMEM16F CaPLSases, and regulates trophoblast fusion.

Taken together, our current study discovered a hitherto under-appreciated Ca^2+^-permeable channel, TRPV4 in human trophoblasts and demonstrated that microdomain-coupled Ca^2+^ entry through TRPV4 directly activates TMEM16F CaPLSase and contributes to trophoblast fusion (Fig. 6E).

## Discussion

Compared to the extensively studied activation mechanisms for TMEM16A/TMEM16B CaCCs, the physiological activation mechanisms for TMEM16F remain unclear. This greatly hindered our understanding of TMEM16F’s biological functions in a wide range of physiological and pathological processes, including blood coagulation, bone mineralization, HIV infection, trophoblast fusion and SARS-CoV2-mediated syncytialization. Owning to their higher Ca^2+^ sensitivity, the TMEM16-CaCCs can be readily activated by Ca^2+^ entry and Ca^2+^ release from internal stores in both excitable and non-excitable cells (20–23). However, the reported CaCC activation mechanisms may not be directly applicable to TMEM16F CaPLSase due to its relatively lower Ca^2+^ sensitivity (8, 9, 25). In this study, we used human trophoblasts as a model system and showed that Ca^2+^ entry through TRPV4 is critical for TMEM16F CaPLSase activation under physiological conditions.

Utilizing the specific TRPV4 agonist GSK101, fluorescence imaging, patch clamp recording in the presence of different Ca^2+^ chelators, we showed that TRPV4 and TMEM16F are functionally coupled within local Ca^2+^ microdomains in human trophoblasts. The spatial proximity between the Ca^2+^ channels and TMEM16F ensures higher and more sustained local Ca^2+^ elevation, which is required for TMEM16F activation. Future studies are needed to examine if functional coupling between Ca^2+^ permeable channels and TMEM16F is a generalized mechanism for TMEM16F activation in different cell types. Ca^2+^ permeable channels other than TRPV4 may be identified as Ca^2+^ sources for TMEM16F activation. Given the relatively transient nature of Ca^2+^ release from internal Ca^2+^ stores, Ca^2+^ release might not play a major role in activating TMEM16F in the plasma membrane, another worthy concept to be elucidated in future studies.

As a polymodal temperature, chemical and osmotic sensors in vertebrates, TRPV4 Ca^2+^ permeable channel is widely expressed and has been reported to be important for vascular function, joint biology and disease, nociception, itch, central nervous system regulation of systemic tonicity, neuroprotection, skin barrier, immune and neurosensory function, lung fibrosis, and skeletal integrity (54). TRPV4’s importance in reproduction was first demonstrated in an elegant study of the uterus, in which Ying et al demonstrated that TRPV4 in myometrium modulates uterine tone during pregnancy and contributes to preterm birth (55). However, TRPV4’s expression and physiological function in the human placenta remain unexplored. In our present study, we demonstrated for the first time that TRPV4 is functionally expressed in human trophoblasts and plays a key role in regulating trophoblast Ca^2+^ signaling, activating TMEM16F and subsequently mediating trophoblast fusion (Fig. 6E).

Our discovery of TRPV4 expression in human trophoblasts also opens a new avenue to further understand TRPV4 in reproductive biology and pregnancy complications. Future studies are needed to understand how TRPV4 is activated under physiological and pathological conditions and if it participates in other trophoblast functions besides activating TMEM16F (Fig. 6E). As a polymodal sensor, TRPV4 senses temperature, osmolarity, and it can be activated by endogenous signaling molecules including bioactive lipid, such as glycerophospholipids (56) and endocannabinoids (54). It is plausible that TRPV4 may sense physiological cues, and respond to changing concentrations of endogenous lipid mediators, to maintain trophoblasts’ functions. On the other hand, inappropriate activation mechanisms of TRPV4 might contribute to some pregnancy complications. Moreover, some of the dominant gain-of-function (GOF) mutations of *TRPV4* are lethal, up to this point considered as critically-severe skeletal malformations. Our data suggest perhaps a co-contribution of gain-of-function of TRPV4 to placental malfunction that in turn contributes to perinatal lethality(57, 58), another interesting subject for future studies.

Cell fusion including cell-cell fusion and viral-cell fusion requires synergistic actions of multiple fusion machineries and is tightly regulated (59–61). In addition to the known fusogens, the recent studies from our and other laboratories showed that TMEM16F CaPLSase also plays essential roles in trophoblast fusion (12), HIV-host cell fusion (14), and SARS-CoV2-mediated syncytialization (13). Although the detailed mechanism for TMEM16F-mediated cell-cell fusion remains to be determined, it is plausible that PS exposure mediated by TMEM16F CaPLSase serves as a ‘fuse-me’ signal to prime the fusogenic sites or fusogenic synapses to facilitate these cell fusion events. Our current finding that TMEM16F is functionally coupled to TRPV4 Ca^2+^ permeable channels raises the provocative question of molecular identity of the Ca^2+^ source that activates TMEM16F in important viral diseases where TMEM16F syncytium-formation plays a significant role, namely HIV infection (14) and SARS-CoV2-mediated pathological syncytialization (13). Targeting the functional coupling between the Ca^2+^-permeable channels (62) and TMEM16F might be a complementary new therapeutic strategy for AIDS and COVID19.

## Materials and Methods

### Cell lines

The BeWo cell line was a gift from Dr. Sallie Permar at Duke University and was authenticated by the Duke University DNA Analysis Facility. The TMEM16F KO BeWo cell line was generated by sgRNAs targeting exon 2 as described previously (12). The HEK293T cells are authenticated by Duke Cell Culture Facility. BeWo cells were cultured in Dulbecco’s Modified Eagle Medium-Hams F12 (DMEM/F12) medium (Gibco, REF 11320-033) and HEK293T cells were cultured in Dulbecco’s modified Eagle medium (Gibco, REF 11995-065). Both media were supplemented with 10% FBS (Sigma, cat. F2442) and 1% penicillin/streptomycin (Gibco, REF 15-140-122) and cells were cultured in a 5% CO_2_-95% air incubator at 37°C.

### Human placenta tissue and primary cultured human trophoblast cells

Placental tissues were collected under the Institutional Review Board approval (IRB# PRO00014627 and XHEC-C-2018-089). Placental cytotrophoblast cells from human term placenta were prepared using a modified method of Kliman as described previously (12). The cytotrophoblasts were plated in fibronectin coated cover glass and supplied with DMEM/F12 medium, 10% FBS, and 1% penicillin/streptomycin.

### siRNA transfection

BeWo cells were seeded on coverslips coated with poly-L-lysine (Sigma, # P2636). One day after seeding, BeWo cells were transfected with TRPV4 siRNAs (Horizonz, # MQ-004195-00-0002) using Lipofectamine RNAiMAX Transfection Reagent (Invitrogen, # 13778075) following the manufacturer’s instructions. After siRNA transfection for 24 hours, the medium was changed to fresh DMEM-F12 and the cells were cultured for another 24 hours before confocal imaging.

### Fluorescence imaging of Ca^2+^ and PS exposure

Ca^2+^ dynamics were monitored using a calcium indicator Calbryte 520 AM (AAT Bioquest, # 20701). BeWo cells were stained with 1 μM Calbryte 520 AM in DMEM-F12 for 15 minutes at 37°C and 5% CO_2_. PS exposure was detected using CF 594–tagged Annexin V (AnV, Biotium, # 29011). To monitor Ca^2+^ and PS dynamics before stimulation, Calbryte 520 AM-stained cells were loaded on the cover glass and incubated in AnV (1:175 diluted in Hank’s balanced salt solution) and image for 50 seconds. To stimulate TRPV4 activity, 20 nM TRPV4 agonist GSK101 (Sigma, #G0798) was used. To inhibit TRPV channel activities, 500 nM TRPV4 antagonist GSK219 (Sigma, #SML0694) was applied to trophoblasts. Cells were continued to be imaged for another 10 minutes. Zeiss 780 inverted confocal microscope was used to image Ca^2+^ and PS dynamics at a 5-second interval. MATLAB was used to quantify the cytosolic calcium and AnV binding.

### BeWo cell fusion and quantification of fusion index

After seeding cells for 24 hours, BeWo cells were treated with 30 μM forskolin (Cell Signaling Technology, #3828s) for 48 hours to induce cell fusion, with forskolin-containing media changed every 24 hours. After forskolin treatment, cells were stained with Hoechst (Invitrogen™, #H3570, 1:2000) and Di-8-ANEPPS (Biotium, #61012, 2 µM) or Wheat Germ Agglutinin Alexa Fluor™-488 Conjugate (Invitrogen™, #W11261, 1:1000) for 15 min at 37°C and 5% CO_2_. For each treatment group, six random fields of view were acquired using Zeiss 780 inverted confocal microscope. Cell fusion is quantified by calculating the fusion index (FI) as described previously (12). FI is calculated as:

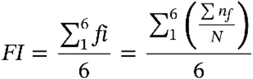

where *fi* represents the fusion index of each field; *nf*, the number of fused nuclei in each field; *N*, the total nuclei number in each field. To avoid the instances of cell division, cells with two nuclei were not considered as fused cells.

### Immunostaining

The mCherry-TMEM16F expressing BeWo TMEM16F knockout (KO) cells were generated by transduction of mCherry-TMEM16F containing lentiviral particles as described previously (12). Flag-tagged R. norvegicus TRPV4 plasmids (Addgene, # 45751) were transiently transfected to the HEK293T cells by using X-tremeGENE360 transfection reagent (Millipore-Sigma). Medium was replaced with regular culture medium 4 hours after transfection. Ruthenium red (Sigma, #R2751, 10μM) was add into the culture medium to inhibit TRPV4 activation. Cells were fixed with 1% paraformaldehyde in phosphate-buffered saline (PBS) for 10 minutes, permeabilized with 0.1% Triton X-100 in PBS, and blocked with 5% goat serum in PBS for an hour. Coverslips were incubated in primary antibodies, anti-TRPV4 (1:200, Alomone, #ACC-034) and anti-mCherry (1:500, NOVUS, #NBP1-96752) or anti-Flag (1:1000, Sigma, #F1804), at 4 °C overnight. Secondary antibodies, Alexa Fluor 488 or Alexa Fluor 640 fluorescence system (Molecular Probes, #35552), were used for fluorescent staining. After staining nuclei with 4′,6-diamidino-2-phenylindole (DAPI), coverslips were mounted using ProLong™ Diamond Antifade Mountant (Invitrogen, #P36961) and imaged with Zeiss 780 inverted confocal microscope.

### Electrophysiology

All currents were recorded in either outside-out or whole-cell configurations on BeWo cells using an Axopatch 200B amplifier (Molecular Devices) and the pClamp software package (Molecular Devices). The glass pipettes were pulled from borosilicate capillaries (Sutter Instruments) and fire-polished using a microforge (Narishge) to reach resistance of 2–3 MΩ. The pipette solution contained (in mM): 140 CsCl, 10 HEPES, 1 MgCl_2_, adjusted to pH 7.2 (CsOH), 0.2 or 2 mM EGTA or 2mM BAPTA as indicated. The bath solution contained 140 CsCl, 10 HEPES, 0 or 2.5 CaCl_2_ as indicated, adjusted to pH 7.4 (CsOH). Pharmacological reagents were applied from extracellular side including 30 nM GSK101 (Sigma, #G0798), 500 nM GSK219 (Sigma, #SML0694) as indicated. Procedures for solution application were as employed previously (5, 26). Briefly, a perfusion manifold with 100–200 μm tip was packed with eight PE10 tubes. Each tube was under separate valve control (ALA-VM8, ALA Scientific Instruments), and solution was applied from only one PE10 tube at a time onto the excised patches or whole-cell clamped cells. All experiments were at room temperature (22–25°C). All the other chemicals for solution preparation were obtained from Sigma-Aldrich.

### Statistical analysis

All statistical analyses were performed with Clampfit 10.7, Excel and Prism software (GraphPad). Two-tailed Student’s t-test was used for single comparisons between two groups (paired or unpaired). One-way ANOVA following by Tukey’s test was used for multiple comparisons. Data were represented as mean ± standard error of the mean (SEM) unless stated otherwise. Symbols *, **, ***, **** and ns denote statistical significance corresponding to p-value <0.05, <0.01, <0.001, <0.0001 and no significance, respectively.

## Supporting information

BeWo With GSK101

TMEM16F KO BeWo With GSK101

## Author contributions

H.Y. and Y.Z. perceived the research and designed the experiments. H.Y. supervised the project. Y.Z. performed all imaging and immunostaining experiments. P.L. performed electrophysiology recording. Z.K.S. performed immune-colocalization experiment. L.F. provided human trophoblast cells and tissues, and performed qRT-PCR measurement. Y.C. and W.L. provided critical reagents and advised on the project. Y.Z., P.L., K.Z.S. and L.F. conducted data analysis. H.Y., Y.Z. and W.L. wrote the manuscript.

## Acknowledgments

This work was supported by NIH-DP2GM126898 grant (to H.Y.). We appreciate Dr. Hua Pan for technical support.

