## Supplementary Information for "Functional coupling between TRPV4 channel and TMEM16F modulates human trophoblast fusion"

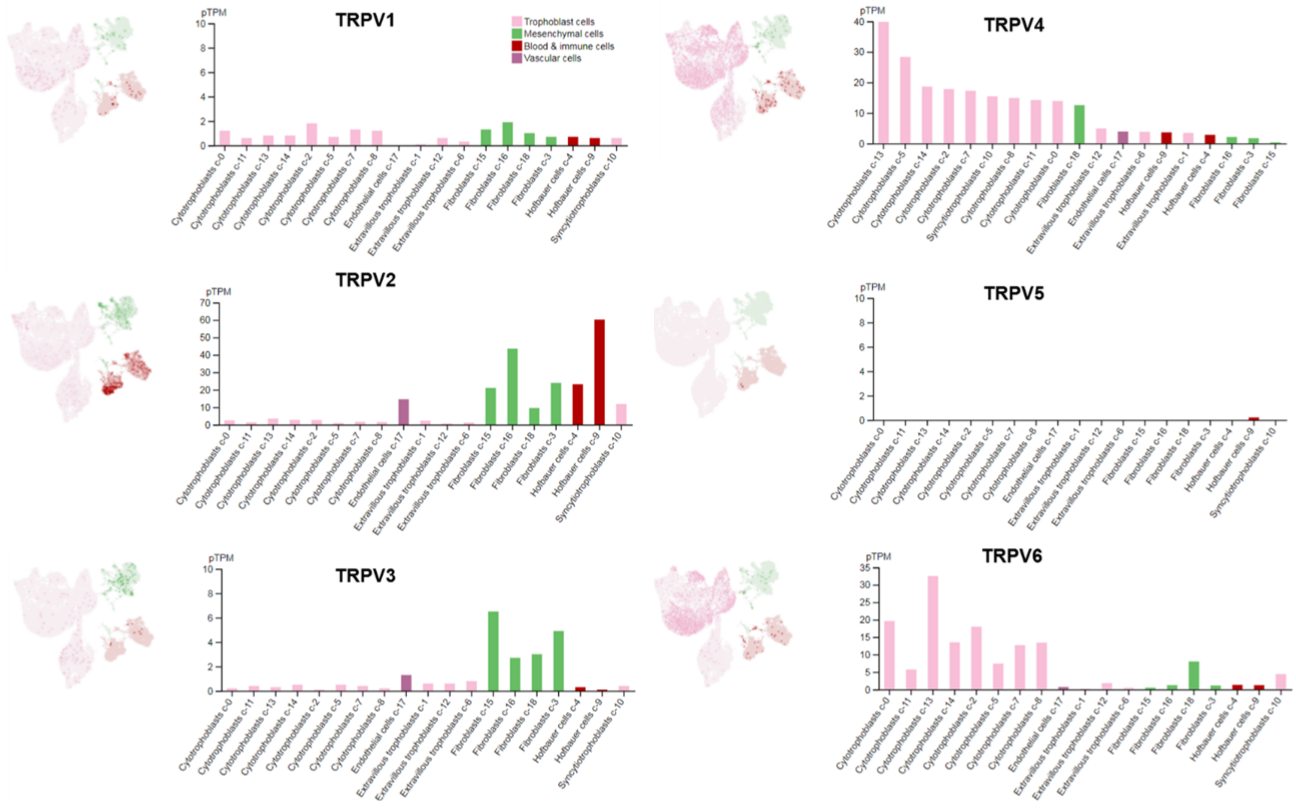


**Figure S1. The single cell RNA sequencing results of TRPV channels in different cell types from human placentas.** The UMAP plots (left) illustrate single cell RNA expression profiles in different cell types, and the bar graphs (right) show pTPM levels in each single cells cluster. All the data were from the Human Protein Atlas V20.1.proteinatlas.org.

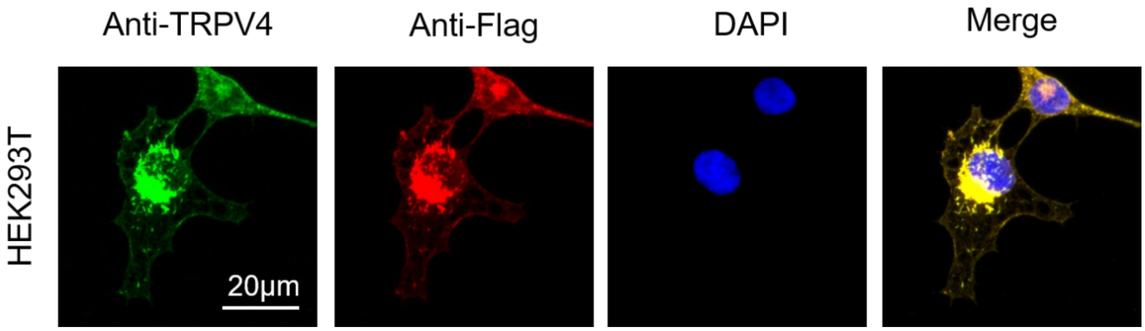


**Figure S2. Validation of the TRPV4 antibody in HEK293T cells heterologously expressed a flag-tagged TRPV4 plasmid.** Immunofluorescence of TRPV4 (anti-TRPV4, green) and Flag tag (anti-Flag, red) in HEK293T cells transfected with a Flag-tagged TRPV4 plasmid. DAPI stained the nuclei and was shown in blue.

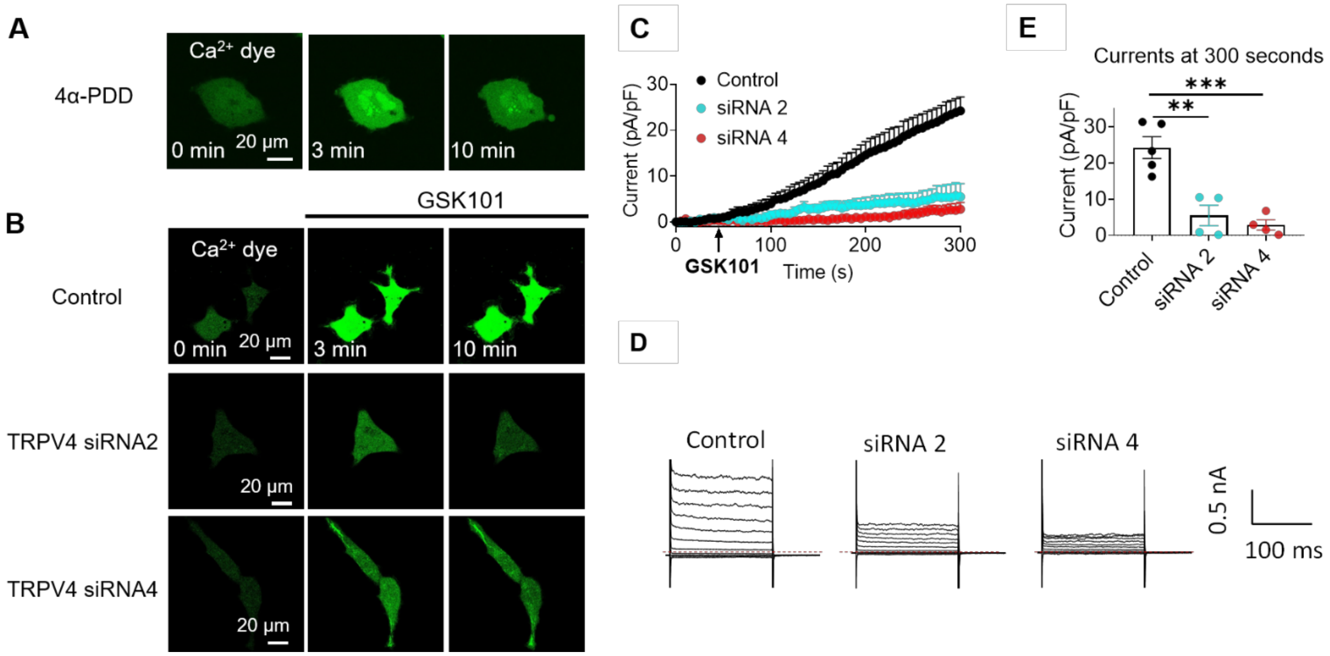


**Figure S3. siRNA knockdown further supports TRPV4 functional expression in trophoblasts. (A)** 4α-phorbol-12,13-didecanoate (4α-PDD, 20 μM), a TRPV4 agonist, triggered intracellular $Ca^{2+}$ elevation in BeWo cells. **(B)** GSK101-induced $Ca^{2+}$ influx diminished in TRPV4 siRNA knockdown BeWo cells. BeWo cells transfected with control siRNA or two different TRPV4 siRNAs (siRNA2 and siRNA4) were stained with Calbryte 590 to visualize cytosolic $Ca^{2+}$ dynamics. **(C)** Representative currents elicited by GSK101 in scrambled-siRNA group and TRPV4 siRNA groups. The experiments were done with whole-cell patch clamp. The currents were elicited with a ramp protocol every 5 seconds and plotted at +100 mV. Error bar represents SEM. **(D)** Representative current traces elicited by a voltage step protocol (200 ms) from -100 mV to +140 mV. **(E)** Currents densities at +100 mV after GSK101 application. Current densities from BeWo cells treated with scrambled siRNA, TRPV4-siRNA2 and TRPV4-siRNA4 were 24.24±3.02, 5.47±2.86 and 24.24±3.02 pA/pF, respectively. Values represent mean ± SEM and statistics were done using one-way ANOVA followed by Tukey's test. (**: p<0.01 and *** : p<0.001, n=4-5).

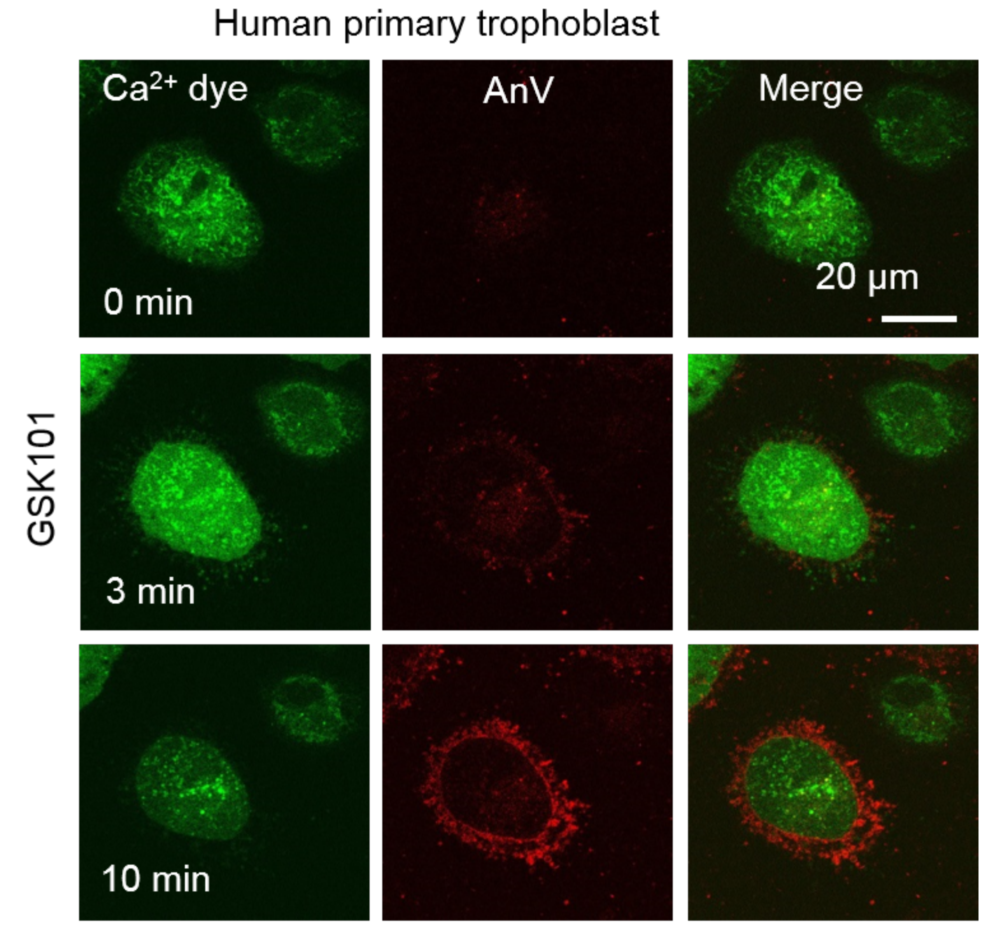


**Figure S4. 20 nM GSK101 triggered $Ca^{2+}$ increase and subsequent CaPLSase activities in primary human placental trophoblasts.** $Ca^{2+}$ dye (Calbryte 520) and fluorescently tagged AnV proteins (AnV-CF594) were used to measure the dynamics of intracellular $Ca^{2+}$ and PS externalization, respectively. All fluorescence images are the representatives of at least three biological replicates.

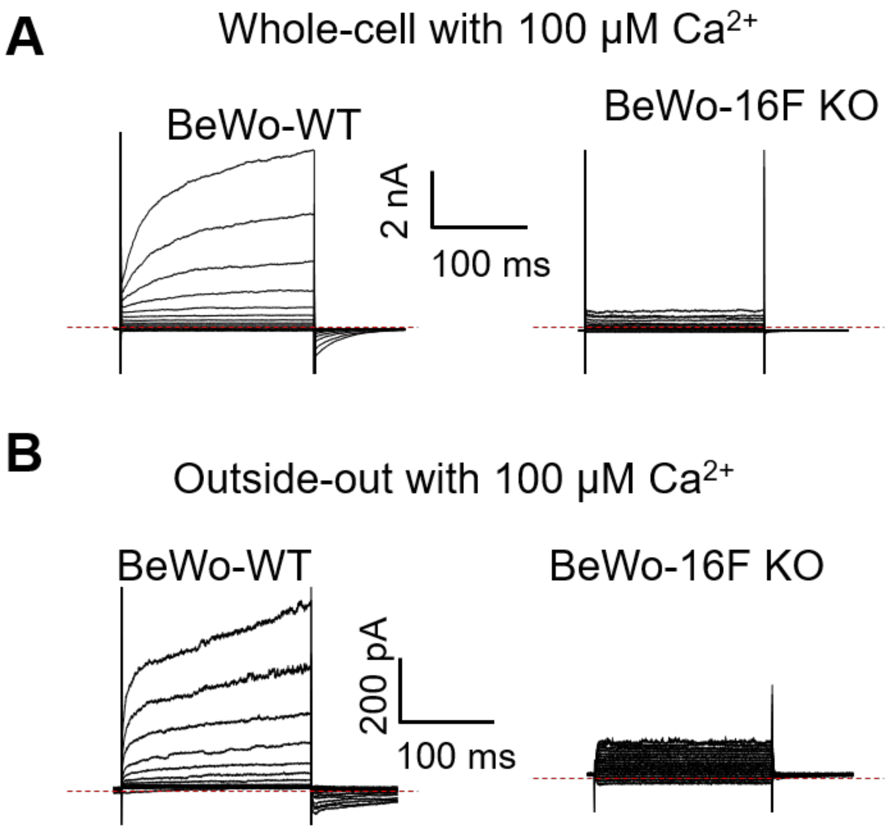


**Figure S5. $Ca^{2+}$-activated current in BeWo wildtype (WT) and TMEM16F knockout (KO) cells. (A)** Representative whole cell recordings elicited by a voltage step protocol (200 ms) from -100 mV to +140 mV. The currents were recorded at ~7-10 minutes after whole-cell patches were formed. The free $Ca^{2+}$ inside the pipette was 100 µM. **(B)** Representative outside-out patch recordings elicited by a voltage step protocol (200 ms) from -100 mV to +140 mV. The currents were recorded immediately after outside-out patches were formed. The free $Ca^{2+}$ inside the pipette was 100 µM.

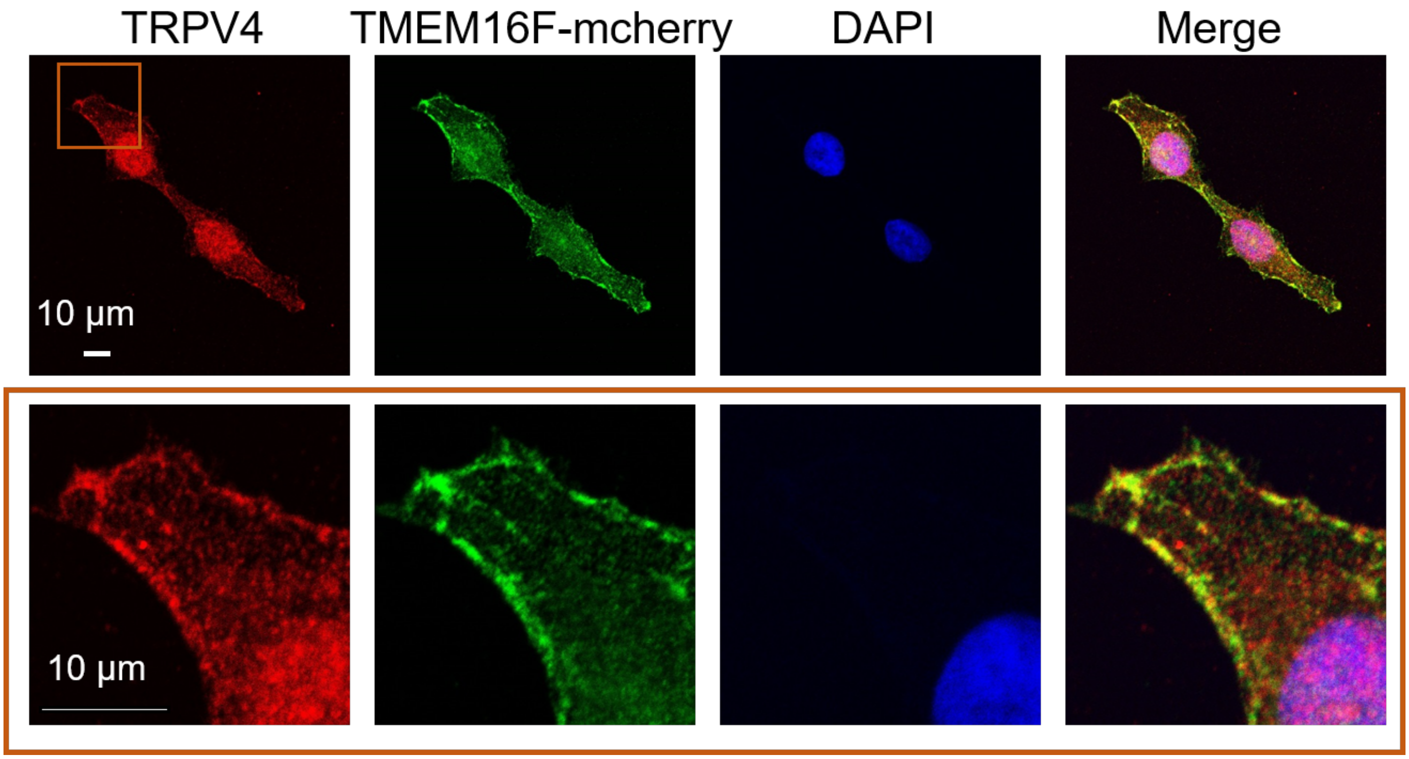


**Figure S6. TRPV4 and TMEM16F are co-localized on BeWo cell membrane.** (Top) Immunofluorescence of the endogenous TRPV4 (anti-TRPV4, red) and the heterologously expressed mCherry-tagged TMEM16F (anti-mCherry) in BeWo TMEM16F knockout cells. (Bottom) The enlarge views of the red box region.

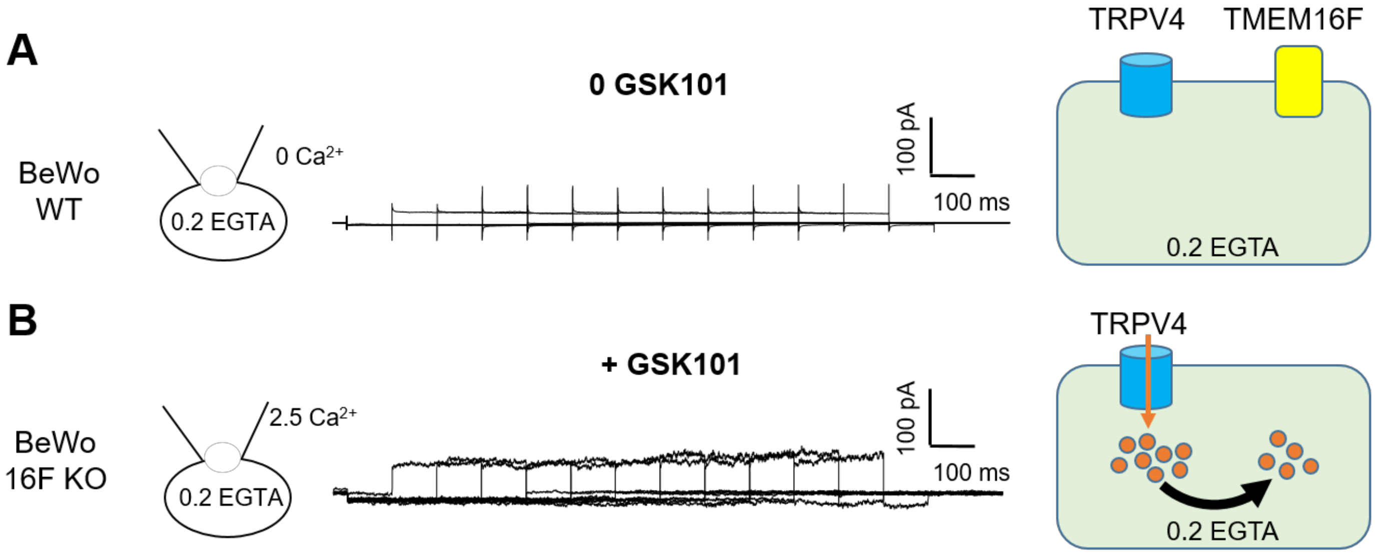


**Figure S7. TRPV4-TMEM16F coupling requires $Ca^{2+}$ influx and TMEM16F expression. (A)** TRPV4-TMEM16F coupling failed to establish when TRPV4 was not activated in wildtype (WT) BeWo cells (n=5). **(B)** 30 nM GSK101-induced TRPV4 activation and $Ca^{2+}$ influx failed to elicit TMEM16F current in the TMEM16F knockout (KO) BeWo cells (n=5). The experimental protocol was the same as shown in Figure 5A.

**Caption for Video S1:** Time lapse video showing wildtype BeWo cells in response to 20 nM GSK101 stimulation. $Ca^{2+}$ dye (Calbryte 520, green) and fluorescently tagged AnV proteins (AnV-CF594, red) were used to monitor the dynamics of intracellular $Ca^{2+}$ and PS externalization, respectively. Related to Fig. 2A.

**Caption for Video S2:** Time lapse video showing TMEM16F knockout (KO) BeWo cells in response to 20 nM GSK101 stimulation. $Ca^{2+}$ dye (Calbryte 520, green) and fluorescently tagged AnV proteins (AnV-CF594, red) were used to monitor the dynamics of intracellular $Ca^{2+}$ and PS externalization, respectively. Related to Fig. 2B.
